# AI-Driven Analysis Unveils Functional Dynamics of Müller Cells in Retinal Autoimmune Inflammation

**DOI:** 10.1101/2025.02.28.640907

**Authors:** Guangpu Shi, Vijayaraj Nagarajan, Rachel R. Caspi

## Abstract

Müller cells are the most abundant glial cell type in the human and mouse retina, playing a crucial role in maintaining retinal homeostasis. However, many aspects of Müller cell function remain poorly characterized. In this study, we reanalyzed a single-cell RNA-seq (scRNA-seq) dataset from Aire-/- mice, focusing on Müller cells and T cells. We identified nine distinct Müller cell subgroups and five T cell subgroups, with activated Müller cells comprising the majority of the Müller cells in the inflamed retina. Using SCassist, an AI-based workflow assistant for single-cell analysis, we created a comparison matrix to quantify pathway involvement in each Müller cell subset. This approach unveils the functional dynamics of Müller cells during retinal inflammation. Activated Müller cells primarily exhibit enhanced inflammatory activity and adopt a macrophage or dendritic-cell-like phenotype, in the presence or absence of increased neuronal activity. These changes are primarily driven by Interferon Regulatory Factors (IRFs), acting alone or in concert with Neuronal Differentiation 1 (NEUROD1). We further inferred the interactions between Müller cells and T cells and found that activated Müller cells do not appear to exhibit extra chemoattraction to Th1 cells compared to other Müller cell subsets, but they do show such effects on the Th1-like regulatory T cells (Tregs). Activated Müller cells display nearly exclusive expression of immune checkpoint molecules, primarily targeting Th1 cells. These findings may uncover a previously unrecognized role for activated Müller cells in attenuating Th1 cell activity.

## Introduction

Müller cells, first described by Heinrich Müller, are the most common type of glial cells in the human and mouse retinas, followed by astroglia and microglia. Müller cells are the only glial cell type that, like retinal neurons, derives from retinal progenitor cells, whereas astrocytes and microglia migrate into the developing retina from external sources. ^1-3^ Müller cells play a crucial role in maintaining retinal homeostasis. Currently, many roles of Müller cells in regulating retinal function remain poorly understood. ^4^ Previous studies have shown that, in addition to providing structural support to the retina ensuring the viability and stability of retinal cells, Müller cells also play several other critical roles. These include supplying trophic substances to neurons, providing antioxidant molecules, removing metabolic waste through phagocytosis, and sustaining synaptic activity via neurotransmitter recycling. ^5-7^ Ions and signaling molecules, including cAMP, are actively involved in these processes. ^4,8^

Retinal inflammation is a central component of human uveitis, a pathogenic autoimmune condition and a major cause of blindness, contributing to 10-15% of severe visual impairment, and affecting particularly working age adults. ^9,10^ The term “uveitis” is used to define a group of eye disorders with intraocular inflammation, encompassing diseases such as sympathetic ophthalmia, birdshot chorioretinopathy, sarcoidosis, Behcet’s disease, and Vogt-Koyanagi-Harada disease. ^11-13^ In chronic retinal inflammation, it has been observed that Müller cells proliferate and physically fill in the gaps left by damaged or degenerated retinal components (reactive gliosis). ^14^ While animal models have provided valuable insights into the pathophysiology of uveitis, the role of retinal Müller cells in responding to and/or modulating the inflammatory process remains unclear. ^15-19^ Current knowledge of Müller cell-T cell interactions is based primarily on our earlier studies, which have demonstrated that Müller cells can inhibit T cell activation, as evidenced by reduced IL-2 receptor expression on T cells and suppressed antigen-induced T cell proliferation following co-culture with Müller cells. ^20-23^ However, the role of Müller cells in retinal inflammation has not been extensively explored in vivo.

Recent studies have shown that Müller cells may interact with T cells by releasing chemokines or expressing cell surface molecules including immune checkpoint molecules, interacting with receptors expressed on T cells. ^18,19^ Yet, these data were collected from bulk Müller cells, lacking subgroup-specific information, which might have overlooked critical details underlying Müller cell activities. Here, we utilized a scRNA-seq dataset from the inflamed retina of an Aire-deficient (Aire-/-) mouse model to investigate the roles of Müller cell subgroups and their interactions with infiltrating T cells. ^18^ Aire is expressed in medullary thymic epithelial cells, where it is responsible for maintaining central tolerance. Aire-/- mouse develops spontaneous and progressive uveoretinitis as part of a multiorgan autoimmune phenotype due to the loss of central tolerance to autoantigens. ^24,25^

To facilitated scRNA-seq data analysis, we developed SCassist (https://github.com/NIH-NEI/SCassist), an AI-powered workflow assistant that leverages large language models (LLMs) such as Google’s Gemini and Meta’s Llama3 to integrate AI-driven insights into the analysis process. In this study, we exploited the information generated by SCassist to construct a comparison matrix that quantifies pathway involvement across Müller cell subgroups. This comparison matrix provides a functional assessment of Müller cell subpopulations and offers insights into their potential roles, enabling us to rapidly and accurately understand the functional dynamics of Müller cells during retinal inflammation, even in the absence of standardized, functionally descriptive nomenclature for Müller cell subsets.

## Material and Methods

### Data download

Single-cell sequencing (scRNA-seq) dataset was downloaded for re-analysis from Gene Expression Omnibus database (GSE132229). ^18^ This dataset includes retinas from four Aire-/- mice and four wild type (WT) controls. The four female Aire-/- mice on a C57BL/6J background include two groups: two 11-week-old mice with grade 2 retinal inflammation and two 16-week-old mice with grade 3 retinal inflammation. The WT controls are age-matched and share the same genetic background. The grade 2 Aire-/- mice contribute 16,884 cells, the grade 3 Aire-/- mice contribute 12,640 cells, and the WT littermate controls contribute 34,672 cells.

### Single cell sequencing data process

The Seurat package (version 5.1.0) was used to perform data quality control and preprocessing following the standard protocol. ^26,27^ After quality controlling to remove outliers in features, counts and mitochondrial gene percentage, the sequencing data were integrated using the Harmony R package (version 1.2.3). ^28^ Data were normalized using the “LogNormalize” method in Seurat with a scaling factor of 10,000. Centering and scaling the data for 2,000 highly variable genes was performed using the “ScaleData” function.

### Dimension reduction, clustering and visualization

As described previously, these steps were performed using Seurat functions. ^29^ Briefly, cells were clustered based on Principal Component Analysis (PCA) using the “RunPCA” function. The first 20 PCs identified (by Elbow method) were used in the “FindNeighbors” (based on k-nearest neighbor (KNN) graphs) and “FindCluster” (Louvain algorithm) functions. “RunTSNE” and “RunUMAP” functions were used with “pca” as the reduction method, to visualize the data. After clustering the retinal cells, Müller cell and T cells were re-clustered using the “subset” function, through combination of the “idents” and “gene expression” arguments. Rlbp1 >0 was used for Müller cells, while Cd3e > 0 was used for T cells. After performing “subset’, we re-ran the standard workflow shown above to cluster Müller cells along with T cells.

### Trajectory analysis and regulon network inference

For trajectory analysis of Müller cells, we utilized the Monocle3 package (version 1.3.7), a widely used computational tool to infer and visualize cell phenotype transition over time. ^30^ Following trajectory analysis, SCENIC was used to identify regulons, including transcription factors (TFs), governing the transition process. ^31^

### Differentially Expressed Gene (DEG) analysis

To identify genes differentially expressed between cell clusters, we used the “FindMarkers” functions in Seurat and set log2FC >1 and p < 0.05 as criteria to define the significant changes in gene expression. Ingenuity Pathway Analysis (IPA) was used to predict the up-stream regulators driving gene expression changes. Gene Set Enrichment Analysis (GSEA) was performed using ClusterProfiler. ^32^

### Characterizing Müller cell subpopulations using SCassist

To gain insight into the features of each Müller cell subpopulation, we employed SCassist, a home-developed AI based tool. It takes the Seurat single cell object and related outputs, generates relevant data metrics, combines it with the template prompt, submits the augmented prompt to the LLM, parses the LLM’s response, displays the results or save them in a file or appends them to the Seurat object. Based on SCassist’s recommendation, we selected ten pathways with the highest percentage of contributing cells to construct a comparison matrix. It contains module scores for each selected pathway in each Müller cell subgroup, quantifying their involvement within each subgroup. The module scores were calculated using a R package UCell (version 2.8.0). ^33^

### Inferring interactions between Müller cells and T cells

To investigate the interactions between Müller cell subpopulations and T cell subpopulations, we utilized the R package CellChat (version 2.1.2), which calculates the potential interaction scores and the strength of cell–cell communications largely based on the expression levels of ligands and receptors. ^34^ This analysis enabled us to identify the most significant ligand-receptor pairs between Müller cells and T cells, revealing the potential molecular mechanism underlying Müller cell-T cell interaction.

### Statistical analysis

All statistical analyses were performed using the algorithms associated with the R packages used, primarily the Wilcoxon Rank Sum Test for ‘FindMarkers’ in Seurat and UCell, Fisher’s exact test and hypergeometric test for ClusterProfiler, and Random Forest Regression and Benjamini-Hochberg test for SCENIC.

## Results

### Müller cells undergo significant shifts in subpopulation proportions during retinal inflammation

Re-clustering of the Müller and T cells from the inflamed retinas and WT controls identified nine distinct Müller cell subpopulations alongside five T cell subpopulations, comprising a total of 1,336 Müller cells and 568 T cells (Figure 1A). As predicted, we noticed that the inflammation increases the diversity of Müller cells, leading to an expansion of the Müller cell subgroups from seven in WT to nine in Aire-/- retina, with the emergence of two new subgroups, Mu_1 and Mu_2 (Figure 1, A, C and D). The proportion test reveals a significant increase in subgroup proportions of Mu_1, Mu_2, Mu_4, and Mu_5 in the Aire-/- retina compared to WT, along with a significant decrease in Mu_8, Mu_7, Mu_6, and Mu_9, in that order (Figure 1, C and D). In contrast, Mu_3 does not exhibit significant changes in its proportion. A cross-condition comparison performed using Seurat did not reveal any significant changes in gene expression either (Mu_3 of Aire-/- vs Mu_3 of WT). In WT Müller cells, Mu_8 alone accounts for over 60% of the cell numbers, while in the Aire-/- retina, the top three subgroups exhibiting the most pronounced increase in proportions (Mu_1, Mu_2, and Mu_4) comprise over 60% of the total Müller cell population. Notably, Mu_1, Mu_2 and Mu_4 are characterized by expression of Gfap, Tead1 and Nfkb1, which serve as markers for mouse Müller cell activation (Figure 1E). ^35^ The log2FD values for Mu_1 and Mu_2 reach the maximum limit of the x-axis in proportion test, indicating that these cell subpopulations are newly emerged, while Mu_4, although present as a very small group of cells in WT retina, became considerably expanded (Figure 1, A, C and D).

**Figure 1.**
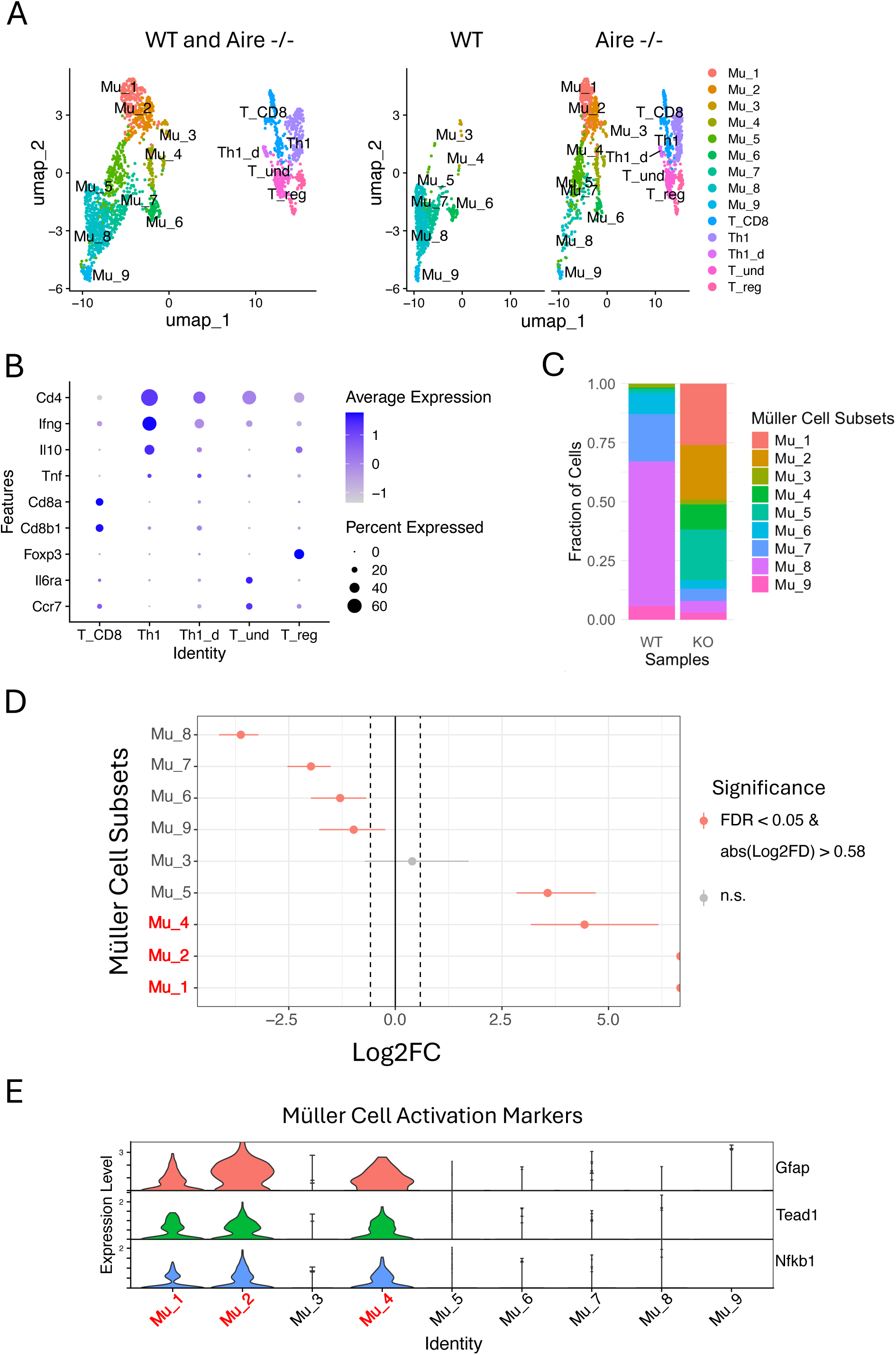
Re-clustering Müller and T cells from Aire-/- retina exhibits significant changes in Müller cell subgroup proportions. (A) A UMAP plot showing nine subgroups of Müller cells and five subgroups of T cells from Aire-/- retinas, using WT retinas as the control. (B) Seurat dot plot showing expression levels of the marker genes used to group T cells. (C) Proportions of Müller cells subgroups from Aire-/- retinas and WT controls. (D) Proportion test showing statistical significance of proportional changes in eight subgroups of Müller cells compared to WT controls (red: activated Müller cells). (E) Expression of the marker genes in Aire-/- retina to distinguish activated Müller cells from the inactive Müller cells (red: activated Müller cells).

Using the reported gene markers for T cell subpopulations, four T cell subgroups were annotated: Th1, T_CD8, Treg and undifferentiated T cells (T_und). ^18^ A small group of CD4+ T cells lacking specific markers for any other T cell subpopulations, but characterized by moderate expression of Ifng, comparable expression of Tnf, and lower expression of Il10 compared to conventional Th1 cells, was annotated as Il10-deficient Th1 (Th1_d) (Fig1B).

### SCassist maps Müller cell subpopulation functions through pathway activity quantification

To characterize the features of each Müller cell subpopulation in the inflamed retina, Seurat identified over 40,000 overlapped marker genes across the nine subpopulations. After analyzing these genes, SCassist recommended approximately 50 pathways from the hundreds of enriched pathways in KEGG or Gene Ontology featuring each subpopulation. We narrowed this list down to 10 pathways with the highest percentage of contributing cells to construct a comparison matrix. It includes module scores for each pathway in each subpopulation, quantifying the extent of pathway involvement within each subpopulation (Figure 2A). Notably, based on the information provided by SCassist, the pathways are grouped into two functionally distinct sections, aligning well with the underlying biology. The top section highlights the well-known functions of Müller cells, ^4,5^ while the bottom section focuses on their responses to retinal inflammation. ^18,19^ Also as expected, the clustering patterns of the columns shown in the comparative matrix logically correspond to the spatial relationships observed in the UMAP (Figure 1A), despite the vastly different data scales involved (hundreds of genes provided by SCassist for comparison matrix versus thousands of genes used for the UMAP).

**Figure 2.**
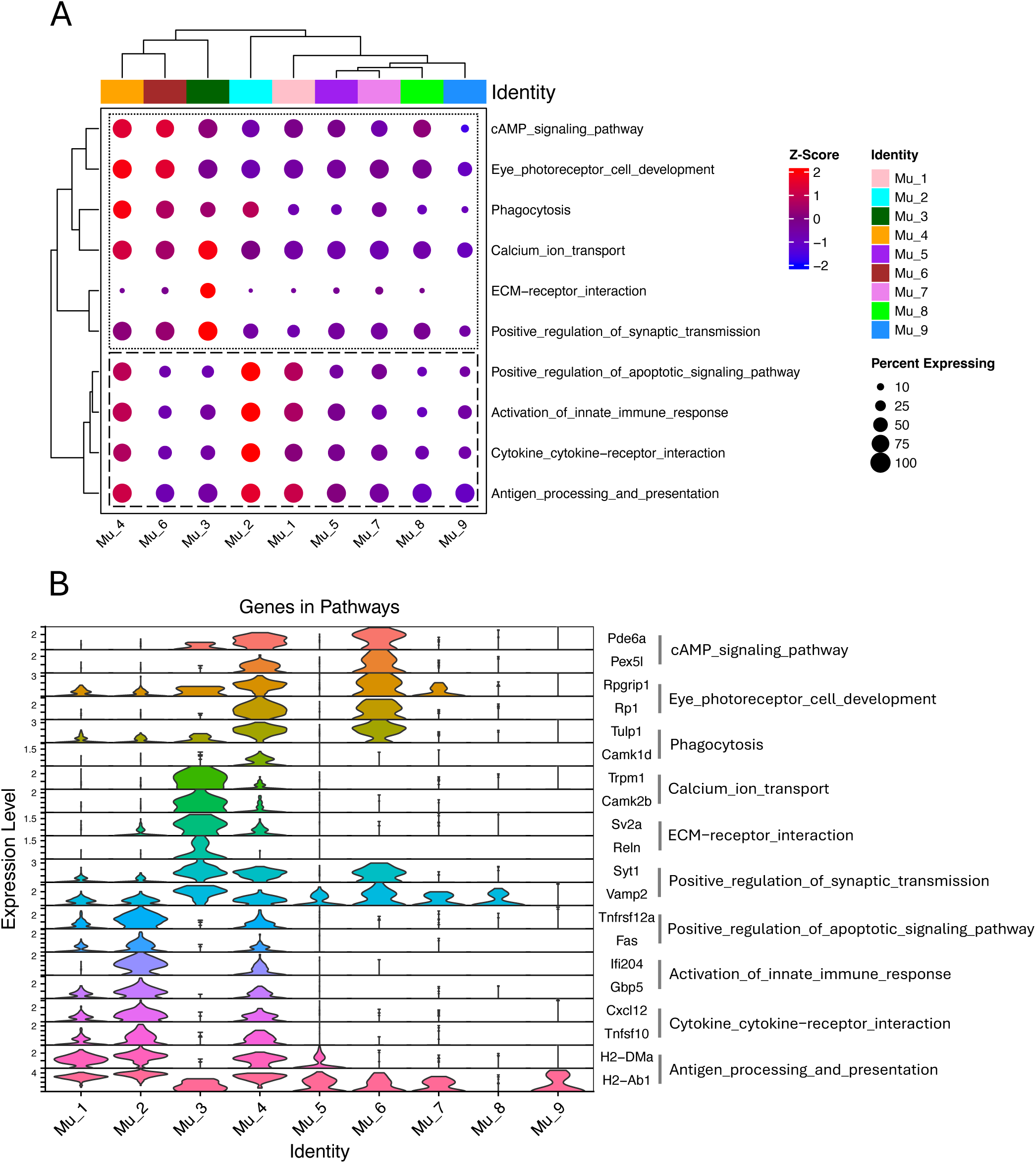
The comparison matrix used to quantify the involvement of pathways in each Müller cell subgroup from Aire-/- retina. (A) The comparison matrix includes module scores for each pathway across subgroups, calculated primarily based on the expression levels of genes associated with the pathways within each subgroup. Top section includes pathways irrelevant to inflammation. Bottom section includes pathways relevant to inflammation. (B) The representative genes used to calculate the module scores for each pathway.

In this comparison matrix, the functional dynamics of Müller cells during retinal inflammation can be appreciated by comparing activated cells (Mu_1, Mu_2 and Mu_4) with inactivated cells (e.g., Mu_8 and Mu_7), which are also found in the WT retina. The activated Müller cell subpopulations exhibit higher module scores than others with Mu_2 being the most prominent for the pathways in the inflammation section, representing the most active response to inflammatory stimuli as reported previously (Figure 2, A and B). ^18-21^ Tnfsf10 and CC/CXC chemokines are the primary contributors to enrichment of the “cytokine-cytokine receptor interactions” pathway, while interferon-induced proteins play a key role in the enrichment of the “innate immune response.” MHC class II molecules and TAP complex components are essential for the enrichment of the “antigen processing and presentation” pathway (Figure 2B). Among the CC and CXC chemokines, we identified Ccl2, CCl27a, Ccl4, Ccl5, Ccl7, Ccl8, Cxcl10 and Cxcl16 in the Aire-/- Müller cell subgroups but only CCl27a, Cxcl10 and Cxcl16 in the WT Müller cells (Figure 6C and Figure S1). Despite the similar trend of module scores in the inflammation section for the activated Müller cells, they behave distinctly in the top section of comparison matrix, where Mu_4 continues to exhibit high module scores, while module scores are lower for Mu_1 and Mu_2.

### Müller cells exhibit a profound phenotype transition in response to retinal inflammation

In the comparison matrix, the activated Müller cells exhibit significant involvement in the activities of antigen presentation, cytokine/chemokine interactions with their receptors, and innate immune response (Figure 2A). This immunological profile is reminiscent of macrophages or dendritic cells. To confirm this phenotype, we performed a cross-condition comparison of gene expression between Müller cells in Aire-/- retina and those in WT control. It revealed that approximately 40% of genes in Aire-/- Müller cells exhibit significant asymmetrical expression changes compared to WT controls. Among them, 781 genes have over two folds upregulation and 30 genes have over two folds downregulation (Figure 3, A and B). Prediction analysis of upstream regulators driving these gene expression changes identified Ifng as the top driving force (Figure 3C). While Th1 cells are the primary source of IFNg, Th1_d cells represent the second most prominent contributors (Figure 1B and Figure 3C, inset panel). To identify the new phenotypes of Aire-/- Müller cells, we conducted GSEA analysis using the cell-type-signature gene sets in MSigDB. The results indicate that Müller cells in the inflamed retina undergo a global phenotypic shift toward a macrophage or dendritic-cell-like phenotype (Figure 3D). The primary genes contributing to the new phenotype are confined in activated Müller cells, indicating that the macrophage or dendritic-cell-like phenotype inferred from GSEA is primarily associated with activated Müller cells, as demonstrated by the comparison matrix (Figure 3, B and E; Figure 2A).

**Figure 3.**
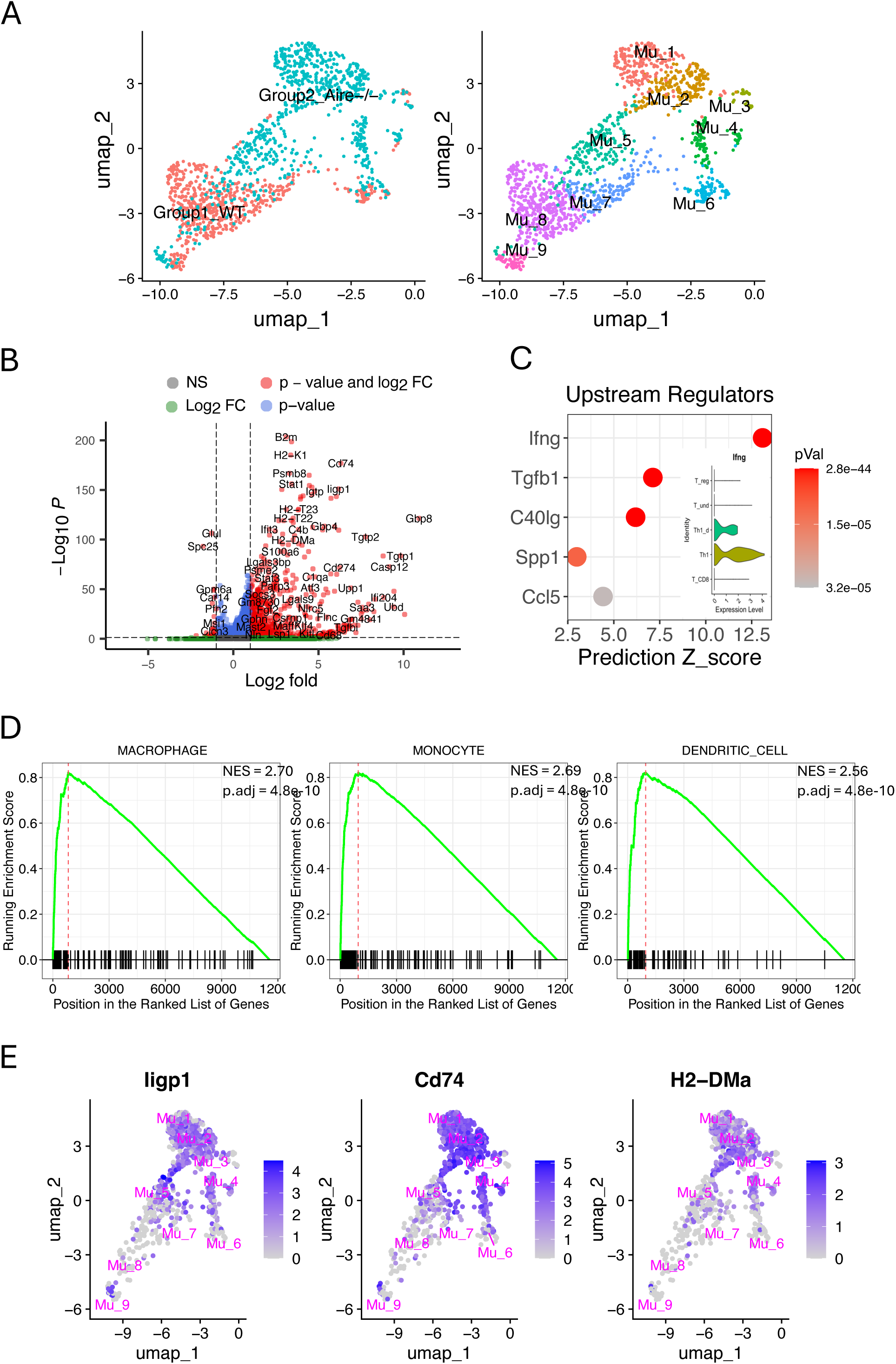
Cross-condition comparison of gene expression levels between Müller cells from Aire-/- retinas and WT controls. (A) Shift of distribution for Müller cells from Aire-/- retina compared to WT control in UMAP. (B) Significant asymmetrical gene expression changes in Aire-/- Müller cells compared to WT controls. (C) Predicted up-stream regulators using IPA, driving gene expression changes in Aire-/- Müller cells, and the sources of IFNg (inset). (D) GSEA analysis supporting the new phenotype of Müller cells in Aire-/- retinas globally. (E) Expression of the important genes significantly upregulated in Figure 3B, which are primarily identified in activated Müller cells.

### Trajectory analysis revealed two transition paths leading to distinct activated Müller cell subpopulations in the inflamed retina

To further explore the details of the phenotypic changes for Müller cells in the inflamed retina, we used Monocle3 to perform a trajectory analysis, tracking phenotypic transition through each population over time. It revealed that Mu_8 cells diverge into two primary branches: trajectory-1 progresses from Mu_8 to Mu_1 and Mu_2 via Mu_5, while trajectory-2 extends from Mu_8 to Mu_4 via M_6 and Mu_7 (Figure 4A). This is consistent with another observation from comparison matrix, where Mu_1 and Mu_2 display similar pattern of module scores, while Mu_6 is grouped with Mu_4 (Figure 2A). Mu_3 may not be involved in phenotype transition, as there are no changes observed in its proportion or gene expression (Figure 1, C and D). Along trajectory-1, Müller cells undergo significant changes in the expression of 606 genes, leading to an activated phenotype for Mu_1 and Mu_2, while, along trajectory-2, Müller cells exhibit significant changes in the expression of 872 genes, resulting in an activated Mu_4 phenotype (Figure 4A and Figure S2). Classification of these genes using Gene Ontology (GO) reveals that the up-regulated genes in trajectory-1 are primarily pertinent to protein synthesis and inflammatory responses, while the up-regulated genes in trajectory-2 include additional genes involved in eye development and light perception, aligning with the information from comparison matrix revealing that Mu_1 and Mu_2 exhibit high scores for inflammation pathways, while Mu_4 show high scores for photoreceptor development as well (Figure 4B and Figure 2A).

**Figure 4.**
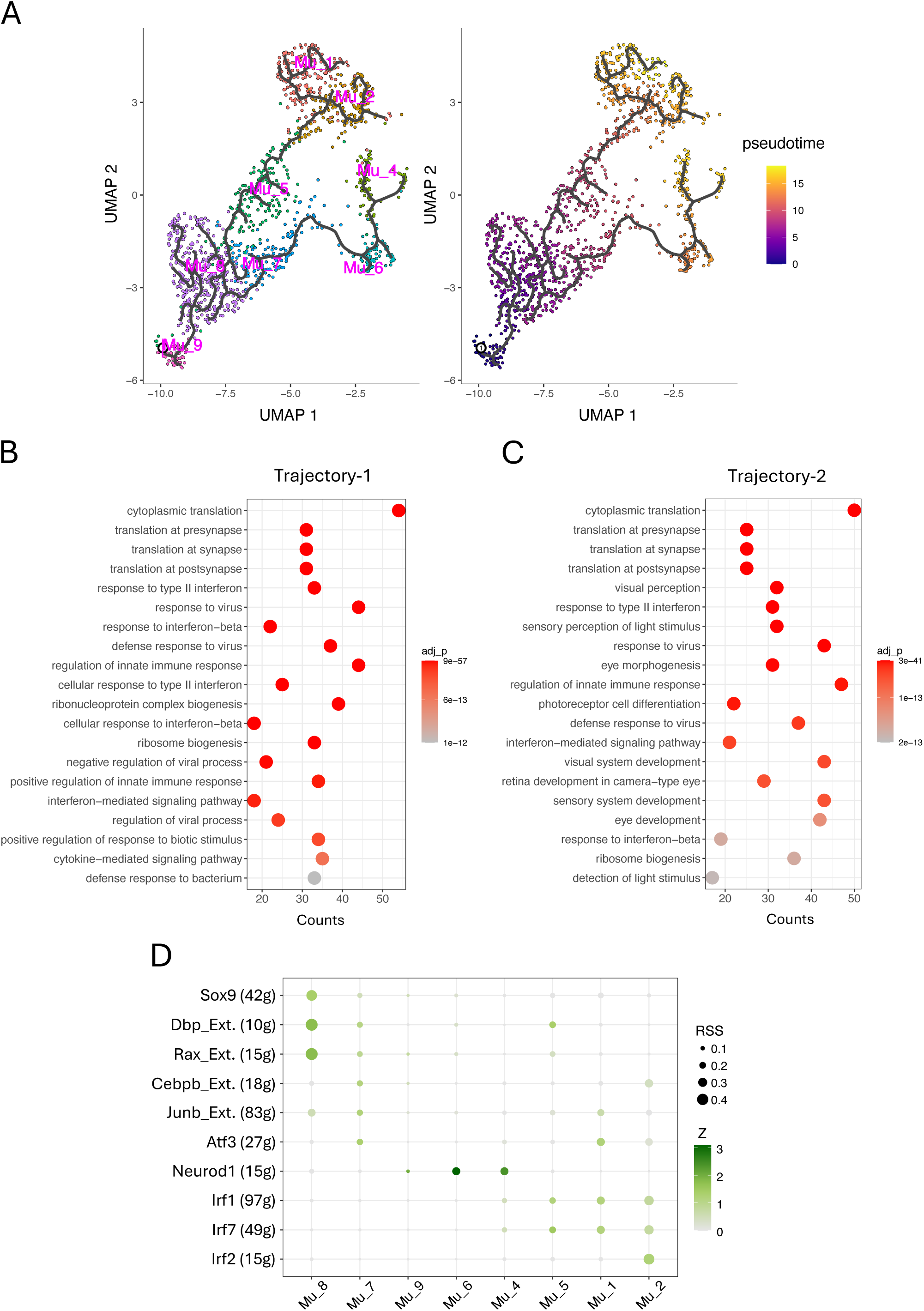
Trajectory analysis of Müller cell phenotypic transition in Aire-/- retina. (A) Two trajectories of Müller cells in Aire-/- retina, leading to activated Müller cells. (B) Top 20 GO terms enriched from genes with significantly upregulated expression along trajectory 1, from Mu_8 to Mu_1 and Mu_2. (C) Top 20 GO terms enriched from genes with significantly upregulated expression along trajectory 2, from Mu_8 to Mu_4. (D) Top 10 regulons inferred by SCENIC, which drive the phenotypic transition of inactive Müller cells along the two trajectories into activated states (Ext.: extended list of co-expressed genes).

We also utilized SCENIC to identify the regulon networks governing the processes along these two trajectories with a focus on the top ten regulons (Figure 4D). In trajectory-1, Mu_5 exhibits enriched activities of Irf1 and Irf7, consistent with the comparison matrix, where Mu_5 begins to display moderate scores in the inflammation section (Figure 2A). As the phenotypic transition progresses, Mu_1 displays increased activity of Junb and Atf3, whereas Mu_2 shows elevated activity of Irf2 and Cebpb. Both subsets share high activity of Irf1 and Irf7. In trajectory 2, Mu_6 exhibits enriched Neurod1 activity, aligning with its high module scores in the upper section of the comparison matrix pertinent to neuronal functions such as photoreceptor development, a process regulated by Neurod1 (Figure 2A). ^36^ Following Mu_6, Irf1 and Irf7 take effect, completing the phenotype transition along this trajectory to Mu_4. Neurod1 distinguishes trajectory-2 from trajectory-1.

### Activated Müller cells demonstrate the most extensive interactions with T cells and other Müller cell subpopulations

To explore the novel biological effects of Müller cell subpopulations on the inflamed retina, we first inferred interactions between these cells and infiltrating T cell subpopulations, as well as within Müller or T cell subgroups. A total of 62 categories of ligand-receptor pairs (LR pairs) were identified including Cell-Cell Adhesion, Secreted Signaling and ECM-Receptors according to the annotation by CellChat (Figure 5A). ^34^ Key LR pairs in the Cell-Cell Adhesion category include NCAM, JAM, CADM, ICAM, MHCI/II, Galectin and PD-L1.

**Figure 5.**
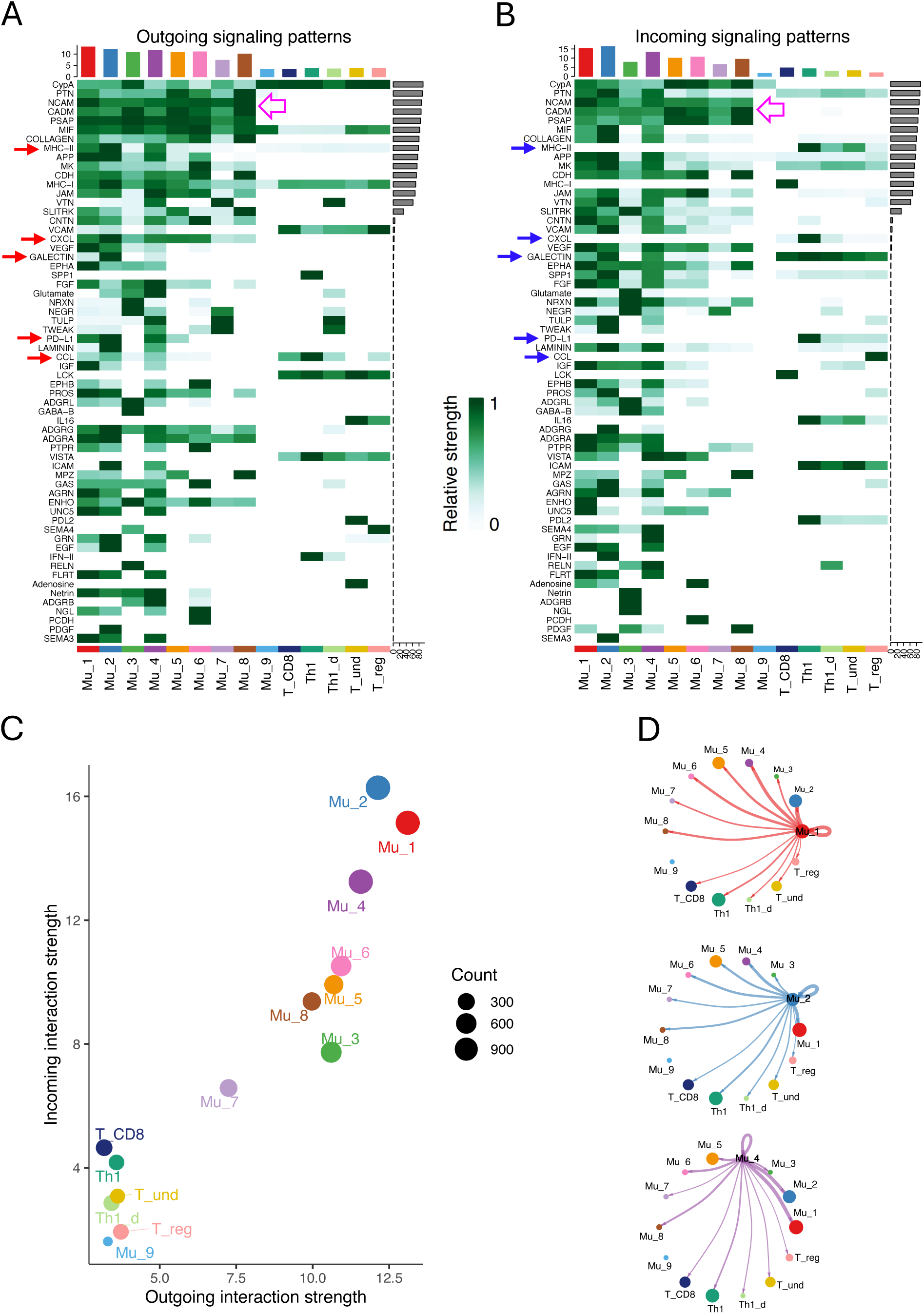
The inferred interactions among Müller cells and T cells in Aire-/- retina. (A) and (B) The sixty-two LR pairs were inferred among all Müller cells and T cells. Red arrows indicate the outgoing signals from the activated Müller cells with the receptor in T cells (blue arrows). Hollow pink arrows specify the interactions between Müller cells, which are mediated primarily by the conventional adhesive molecules. (C) A scatterplot showing the strength of outgoing and incoming signals for all the Müller and T cell subgroups. (D) The circle plots showing interactions between the activated Müller cells and other cell subgroups.

The Secreted Signaling category primarily encompasses cytokines, chemokines, and growth factors.

The three activated Müller cell subpopulations, Mu_1, Mu_2, and Mu_4, exhibit the highest scores for both outgoing and incoming signaling strength, with Mu_2 cells ranking the highest. This suggests that activated Müller cells engage in the most extensive interactions with other cell types, beyond interactions within their own populations (Figure 5, C and D). T cells, in contrast to majority of Müller cells, are notably less active in both sending and receiving signals. Within T cell subgroups, as expected, Th1 and CD8 T cells exhibit the highest activities, while Treg cells are the least active.

We focused primarily on the interactions between activated Müller cells and T cells. Five categories of interactions, including CCL, CXCL, MHC-II, GALECTIN and PD-L1, were identified to have both activated-Müller-cell specific outgoing signals (Figure 5A, red arrows) and corresponding receptor signals in T cells (Figure 5B, blue arrows). Mu_1, Mu_2 and Mu_4 commonly produce Cxcl16, Cxcl10, Ccl5, Ccl8, and Ccl27a (Figure 6, A and C). Among them, Cxcl10, Ccl5 and Ccl8 are relatively specific to activated Müller cells, while Cxcl16 and Ccl27 can be produced from other Müller cells. In addition to these three activated-Müller-cell specific chemokines, Mu_2 cells can also produce Ccl2 and Ccl7, while Mu_4 cells can make Ccl4, at marginal levels. On the recipient side, Th1 and Th1_d cells are exclusively targeted, through Cxcr6, by Cxcl16, whereas other T cell can respond to multiple chemokines. Notably, activated Müller cells do not exhibit extra chemoattraction to Th1 cells compared to inactivated Müller cells, but they do show such effects on the Tregs expressing Cxcr3 and Ccr5, particularly through Mu_2 (Figure 6A). The activated Müller cells not only produce chemokines but also express checkpoint inhibitors, including Cd274 and Lgals9, suppressing the T cells (Figure 6B). Notably, Th1 cells are their predominant targets (Figure 6, B and C). Lgals9 expressed by Mu_2 cells exerts the most potent inhibitory signals to the T cells. The analysis also reveals that the activated Müller cells express higher levels of MHC II molecules, specifically H2-Ab1, H2-Aa, H2-Eb1, H2-DMa, and H2-DMb1, compared to other Müller cell subgroups (Fig. 6D and E).

**Figure 6.**
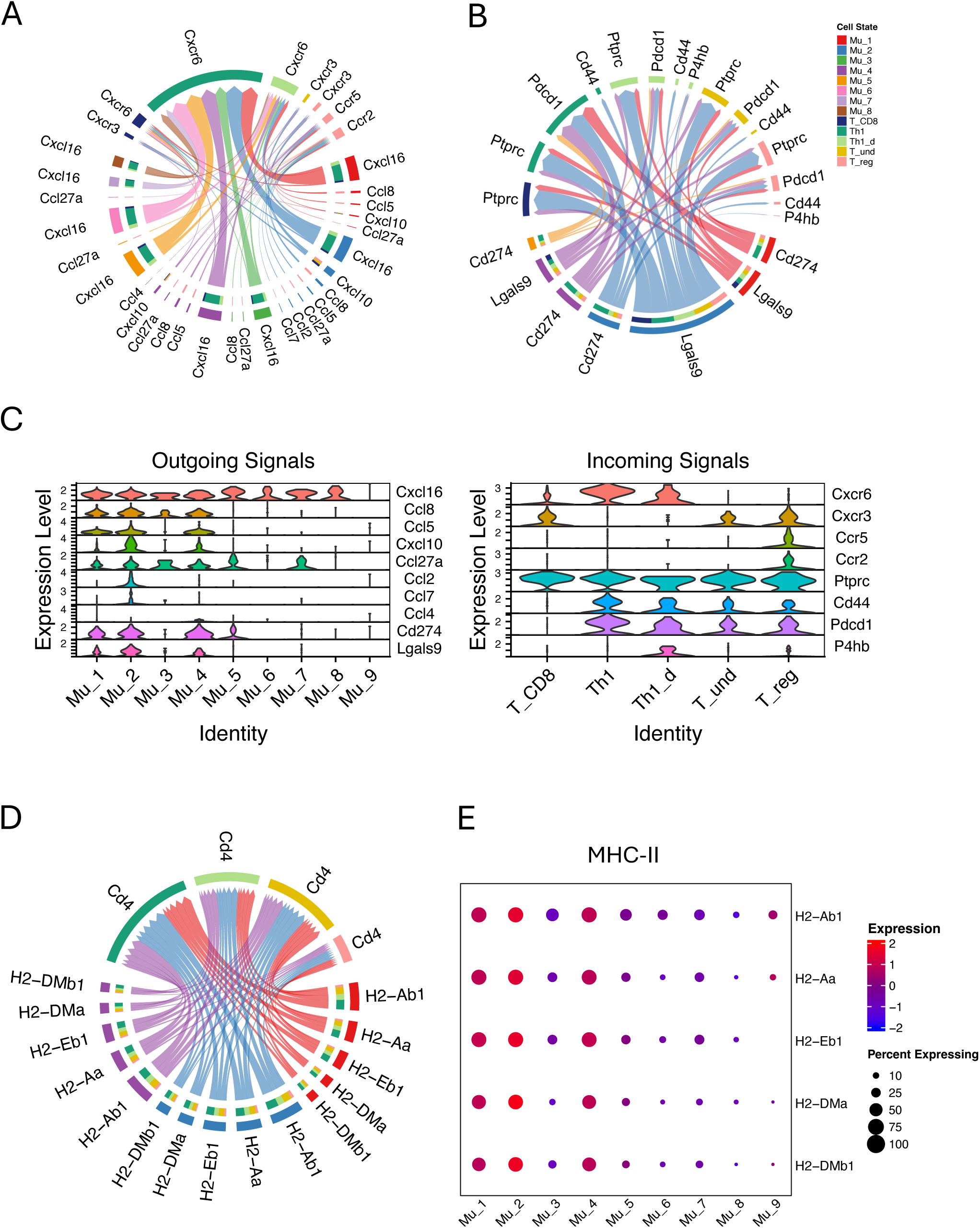
The inferred interactions between Müller cells and T cells through LR pairs of CCL, CXC, PD-L1, Galectin and MHC-II in Aire-/- retina. (A) Müller cells interact with T cells through LR pairs of CCL and CXC. (B) Activated Müller cells interact with T cells through LR pairs of PD-L1 and Galectin. (C) Seurat violin plots showing expression levels of the genes mediating outgoing signals from Müller cells or incoming signals to T cells. (D) Activated Müller cells interact with T cells through LR pairs of MHC-II. (E) A dot plot showing the expression levels of MHC-II molecules in Müller cell subgroups.

## Discussion

This study is the first to use scRNA-seq characterizing distinct subpopulations of Müller cells in the retina with autoimmune inflammation. We identified nine subgroups of Müller cells in Aire -/- retinas, including three activated subtypes. Using our in-house AI tool, SCassist, we analyzed these subsets and revealed macrophage or dendritic-cell-like properties in activated Müller cells, one of which exhibited pronounced neuronal activity. We found that transcription factors primarily from the IRF family, along with NEUROD1, orchestrate the transition of Müller cells from an inactivated to an activated state. Our analysis also identified extensive interactions between activated Müller cells and infiltrating T cells. In contrast to previous reports, activated Müller cells do not exhibit enhanced activity contributing to retinal inflammation.

### Activated Müller cells predominate among Müller cells in the inflamed retina

In WT retina, seven distinct Müller cell subpopulations were identified (Figure 1, A and C). Notably, most Müller cells are concentrated in the Mu_7 and Mu_8 subgroups, consistent with prior studies indicating that mouse Müller cells exhibit relative homogeneity under normal conditions. ^35,37,38^ However, retinal inflammation results in significant redistribution of cells across all subgroups except Mu_3, along with the emergence of two novel subpopulations, Mu_1 and Mu_2 (Figure 1, A, C and D). Activated Müller cells (Mu_1, Mu_2 and Mu_4) account for majority of the total Müller cell population in the inflamed retina, implying their central role in sensing and modulating retinal inflammation. In contrast, WT Müller cells contain only a small subset of activated cells, specifically Mu_4 cells, which can expand significantly during inflammation. The potential role of this subset is discussed in the next section.

### SCassist enables rapid and accurate detection of the new phenotypes in activated Müller cells

In scRNA-seq analysis, functional inference of cell subpopulations heavily relies on annotation algorithms and reference databases of cell-type-specific markers. ^39,40^ However, for newly identified cell types, such as the Müller cell subtypes examined in this study, standardized nomenclature and well-defined cell marker profiles are often unavailable. Performing conventional functional analyses based on marker genes, using tools like Seurat or Scanpy, can be time-consuming when applied to multiple cell subgroups. SCassist offers a rapid and accurate solution to this challenge.

As demonstrated in this study, SCassist integrates the information from multiple databases, including the canonical pathways from KEGG, alongside GO terms, thereby enhancing our understanding of the features of Müller cell subpopulations. In the comparison matrix constructed using enriched pathways and associate genes provided by SCassist, combining the KEGG pathways “Antigen processing and presentation” and “Cytokine-cytokine receptor interactions” with the GO term “Activation of innate immune response” reveals macrophage or dendritic cell-like properties in activated Müller cells, a finding further supported by GSEA (Figure 2A, bottom section; Figure 3, D and E). Although previous studies have examined the inflammatory properties of Müller cells in retinal inflammation using GO terms and MSigDB gene sets, insights into pathway involvement, especially those canonical pathways, at the level of individual subgroups are lacking. ^18,19^

In addition to the inflammatory properties, this comparison matrix also demonstrates that Mu_4 cells, approximately 10% of total Müller cells in the inflamed retina, exhibit enhanced activities in pathways associated with neuronal functions, including “eye photoreceptor cell development”, “cAMP signaling”, and “calcium iron transport”. ^4,8,41^ The critical genes contributing to enrichment of these pathways include Pde6a and Pex5l for cAMP signaling; Trpm1 and Camk2b for calcium transport; and Syt1 and Vamp2 for synaptic transmission (Supplemental table). While the precise function of this subset remains unclear, it is reasonable to speculate that these cells are analogous to the Müller cells in zebrafish, which have the capacity to regenerate photoreceptors following retinal injury, an ability that has been lost in mice and humans over the course of evolution. ^35,42^ This speculation is supported by the expression of Neurod1 in this subset, which plays a key role in driving photoreceptor development (Figure 4D). ^36^ This subset provides important insights into the pathophysiology of Müller cells responding to retinal inflammation; however, it was overlooked in previous studies. ^18,19^

### Phenotype transition to activated Müller cell is primarily driven by the coordinated activity of IRFs and NEUROD1

Th1 has been reported to initiate autoimmune retinal inflammation in Aire -/- mice and other animal models. ^16,18,43^ In Aire-/-mice, retinal Müller cells exhibit a globally altered gene expression profile in response to stimulation by IFNg, a key cytokine produced by Th1 cells. ^18^ A similar observation was reported in Müller cells from a mouse model with experimental autoimmune uveitis (EAU). ^19^ However, these studies did not explore the specific molecular mechanisms underlying IFNg-induced activation of Müller cells. In this article, we uncovered that distinct Müller cell subsets respond to IFNg via coordinated gene regulatory networks involving IRFs and NEUROD1, giving rise to three activated Müller cell states.

IRF1 and IRF7 activities are enriched along both trajectories of Müller cell transition, suggesting they are key drivers of these transitions (Figure 4, A and D). IRF2 and CEBPB activities are enriched in the Mu_2, while JUNB and ATF3 activities distinguish the Mu_1 (Figure 4D). Similar to IRFs, CEBPB, JUNB, and ATF3 are all downstream effectors of IFNg signaling. ^44-48^ Trajectory-2 distinguishes itself from trajectory-1 by NEUROD1 in Mu_4 and Mu_6, a key TF for retinal neuronal differentiation. ^36^ The distinct combinatorial patterns of these transcription factors contribute to the phenotypic differences observed among the Müller cell subgroups, particularly in the activated Müller cells, within the comparison matrix (Figure 2A). Notably, some of these transcription factors may compete for binding to the same promoter motifs, for example, IRF2 with IRF1 and IRF7. ^49^ How these transcription factors interact, whether through competition or cooperation, to drive Müller cell phenotypic transitions is largely unknown.

### Activated Müller cells may play an unanticipated role in mitigating Th1 cells

Previous studies on retinal inflammation have suggested that Müller cells may contribute to the inflammatory process through increased expression of chemokines and MHC-II. ^18,19^ Contrary to these hypotheses, we found that activated Müller cells, which make up the majority of the Müller cell population in the inflamed retina, do not exhibit an enhanced ability to chemoattract Th1 cells, compared to other Müller cell populations including those from WT retina. The activated Müller cells interact with Th1 cells exclusively through Cxcl16 and express this chemokine at a level similar to that from other Müller cells (Figure 6, A and C; Figure S1). Cxcl16 appears to be a native chemokine that may play a crucial role in maintaining retinal homeostasis, although its precise function remains to be elucidated. ^50^ We also observed that checkpoint genes, Cd274 and Lgals9, are expressed predominantly in activated Müller cells and Th1 cells have the greatest extent of being affected by these molecules (Figure 6, B and C). Moreover, we observed that activated Müller cells exhibit extra chemoattraction to Tregs by release chemokines including CXCL10, CCL5, CCL8 and CCL27a. These Tregs, which express Cxcr3 and Ccr5, are referred to as Th1-like Tregs and exert potent inhibitory effects on Th1 cells. ^51^

Consistent with previous studies, we observed that the activated Müller cells exhibit higher MHC-II expression than other Müller cells, however, this does not necessarily indicate that these Müller cells can contribute to inflammation by presenting antigens to T cells (Figure 6, D and E). Our previous studies have demonstrated that the ability of Müller cells to activate T cells via antigen presentation is abolished upon contact with T cells, a process mediated by an immunosuppressive molecule expressed on Müller cells. ^20,21^ Furthermore, recent evidence suggests that MHC-II molecules expressed by non-professional APCs may contribute to T cell exhaustion through an unknown mechanism. ^52^ Together, these findings suggest a previously unappreciated role for activated Müller cells in dampening Th1 cell responses.

In summary, our study demonstrates the value of our AI system in advancing scRNA-seq analyses of retinal autoimmune inflammation, providing rapid and precise information on the key pathways and their associated genes. The creation of a comparison matrix proved to be an effective approach for uncovering the distinctive features of cell subgroups, offering deeper insights into their biological significance and guiding more comprehensive data analysis. Using this approach, we successfully detected novel phenotypes of activated Müller cells during retinal inflammation and identified the gene regulatory networks driving their phenotypic transition. The activated Müller cells exhibit previously unrecognized functions, such as utilizing a native chemokine (Cxcl16) to engage with Th1 cells and an enhanced ability to recruit Th1-like regulatory T cells into the inflamed retina.

## Acknowledgment

We thank Dr. Igal Gery for reading the manuscript and providing suggestions, Dr. Wei Li for discussion on the neuronal properties of Müller cell, and the members of Caspi Lab, along with other colleagues in the branch of Laboratory of Immunology of NEI, for offering their feedback regarding data analysis and interpretation.

---

**Figure S1.**
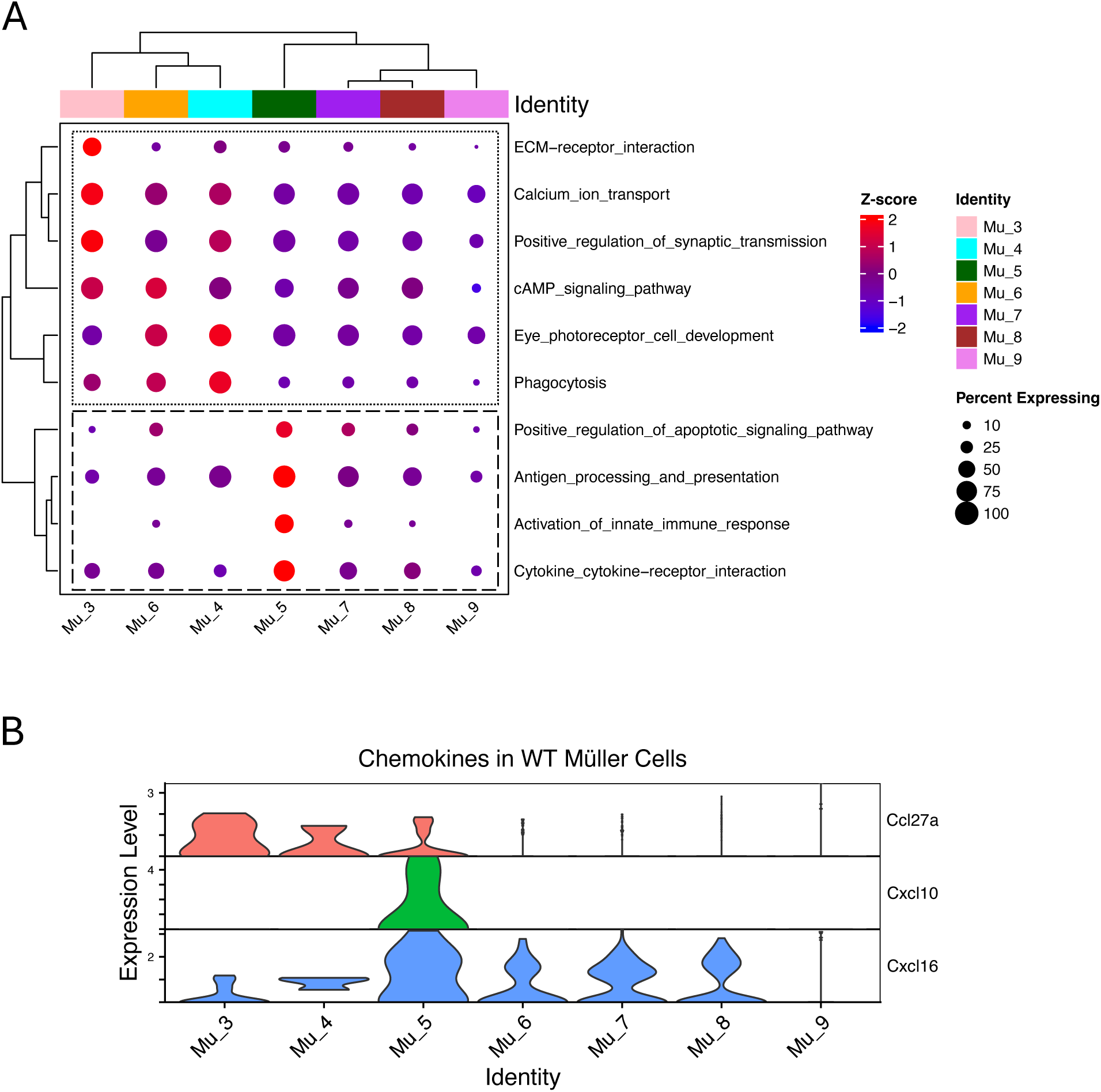
(A) Comparison matrix for the pathways in WT Müller cell subgroups. (B) Chemokine expression in WT Müller cell subgroups.

**Figure S2.**
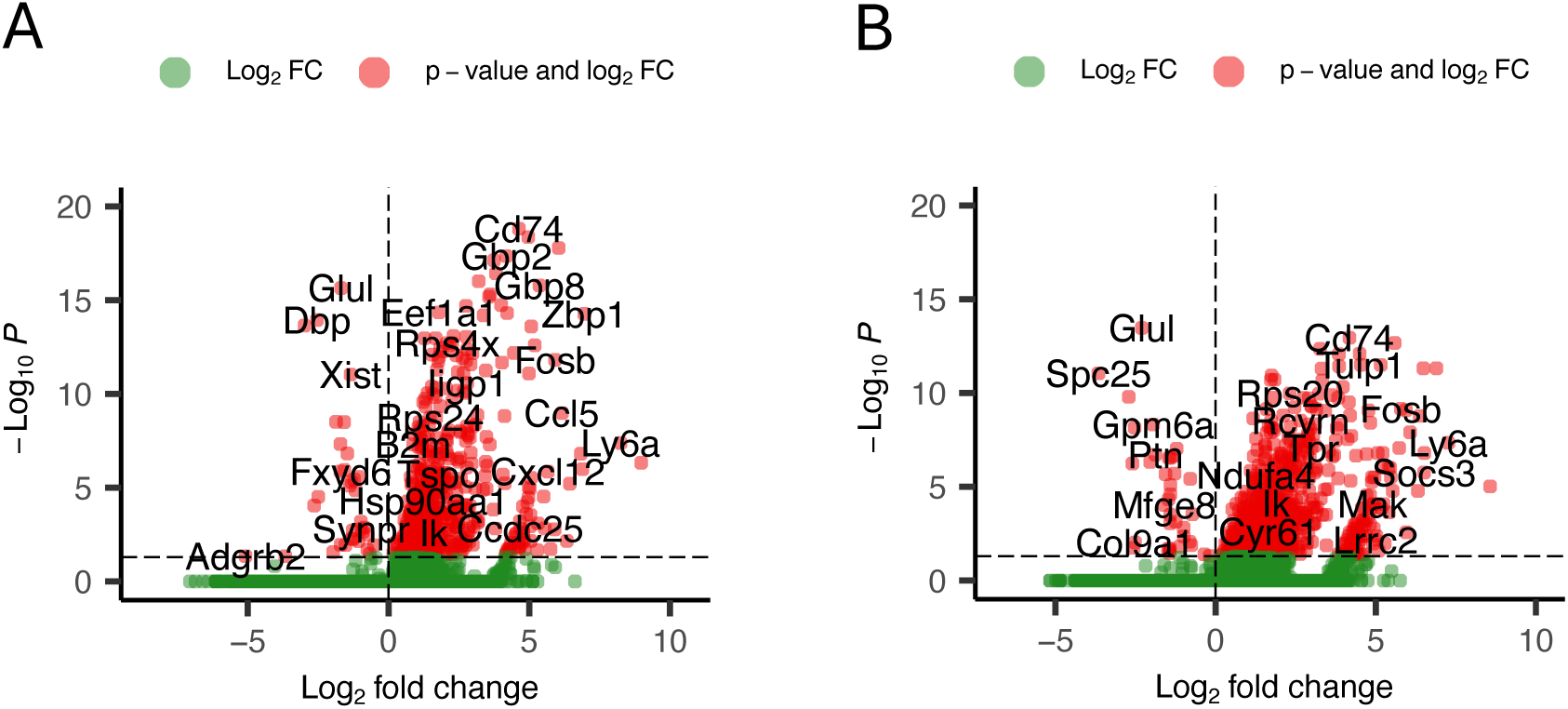
(A) Gene express changes in Trajectory-1. DEGs are identified in Mu_1 and Mu_2 (vs Mu_8). (B) Gene express changes in Trajectory-2. DEGs are identified in Mu_4 (vs Mu_8).

